# Isonitrile Formation by a Non-heme Iron(II)-dependent Oxidase/Decarboxylase

**DOI:** 10.1101/308460

**Authors:** Nicholas C. Harris, David A. Born, Wenlong Cai, Yaobing Huang, Joelle Martin, Ryan Khalaf, Catherine L. Drennan, Wenjun Zhang

**Affiliations:** Department of Chemical and Biomolecular Engineering, University of California Berkeley, Berkeley, California 94720 United States; Chan Zuckerberg Biohub, San Francisco, California, 94158, United States; Department of Plant and Microbial Biology, University of California Berkeley, Berkeley, California 94720 United States; Department of Biology, Massachusetts Institute of Technology, Cambridge, Massachusetts 02139 United States; Graduate Program in Biophysics, Harvard University, Cambridge, Massachusetts 02138 United States; Department of Chemistry, University of California Berkeley, Berkeley, California 94720 United States; Howard Hughes Medical Institute, Massachusetts Institute of Technology, Cambridge, Massachusetts 02139 United States; Department of Chemistry, Massachusetts Institute of Technology, Cambridge, Massachusetts 02139 United States

**Keywords:** Biosynthesis, Acyl-acyl carrier protein ligase, Oxidoreductase, Protein Structures, Isocyanide

## Abstract

The electron-rich isonitrile is an important functionality in bioactive natural products, but its biosynthesis has been restricted to the IsnA family of isonitrile synthases. We here provide the first structural and biochemical evidence of an alternative mechanism for isonitrile formation. ScoE, a putative non-heme iron(II)-dependent enzyme from *Streptomyces coeruleorubidus*, was shown to catalyze the conversion of (*R*)-3-((carboxymethyl)amino)butanoic acid to (*R*)-3-isocyanobutanoic acid through an oxidative decarboxylation mechanism. This work further provides a revised scheme for the biosynthesis of a unique class of isonitrile lipopeptides, members of which are critical for the virulence of pathogenic mycobacteria.

---

The electron-rich functionality of the isonitrile lends itself as a biologically active warhead for naturally derived products. Due to its ability to coordinate transition metals, it is often exploited for metal acquisition, detoxification, and virulence.^1–3^ Indeed the resume of potent biologically active isonitrile containing natural products is vast, and examples include xanthocillin, an antiviral agent;^4^ rhabduscin, a virulence associated phenoloxidase inhibitor;^2^ and many marine sponge derived metabolites (**Figure S1**). Despite the widespread utility of isonitrile in nature, its biosynthesis has long been considered endemic to the IsnA family of isonitrile synthases, which typically convert an α-amino group to isonitrile on an amino acid and require ribose-5-phosphate as a co-substrate.^1,5–8^

Our recent genome mining of a conserved gene cluster widely present in Actinobacteria indicated an alternative route for isonitrile formation.^9^ In particular, we identified and proposed the function of five genes required for the biosynthesis of a unique class of isonitrile lipopeptides (INLPs) that are critical for the virulence of pathogenic mycobacteria. Taking the pathway from *Streptomyces coeruleorubidus* as an example, the biosynthesis was proposed to start with the activation and loading of crotonic acid onto ScoB, an acyl carrier protein (ACP) by ScoC, an acyl-ACP ligase. A Michael addition of Gly to the β-position of crotonyl-ScoB is then promoted by ScoD, a thioesterase to form a Gly adduct **1**, followed by oxidation and decarboxylation, presumably catalyzed by ScoE, a non-heme iron(II)-dependent oxidase, to generate a β-isonitrile fatty acyl-ACP intermediate **2**. This β-isonitrile acyl moiety is then condensed to both amino groups of Lys promoted by ScoA, a single-module non-ribosomal peptide synthetase (NRPS), and reductively released to form a terminal alcohol product **3** (Figure 1). ScoE thus represents a new family of enzymes distinct from isonitrile synthases that promote the transfer of one carbon from ribose-5-phosphate to an amino group to form isonitrile. Although the function of ScoE was reconstituted in *E. coli* for **3** biosynthesis, *in vitro* reconstitution of its activity based on the proposed biosynthetic pathway repeatedly failed.

**Figure 1.**
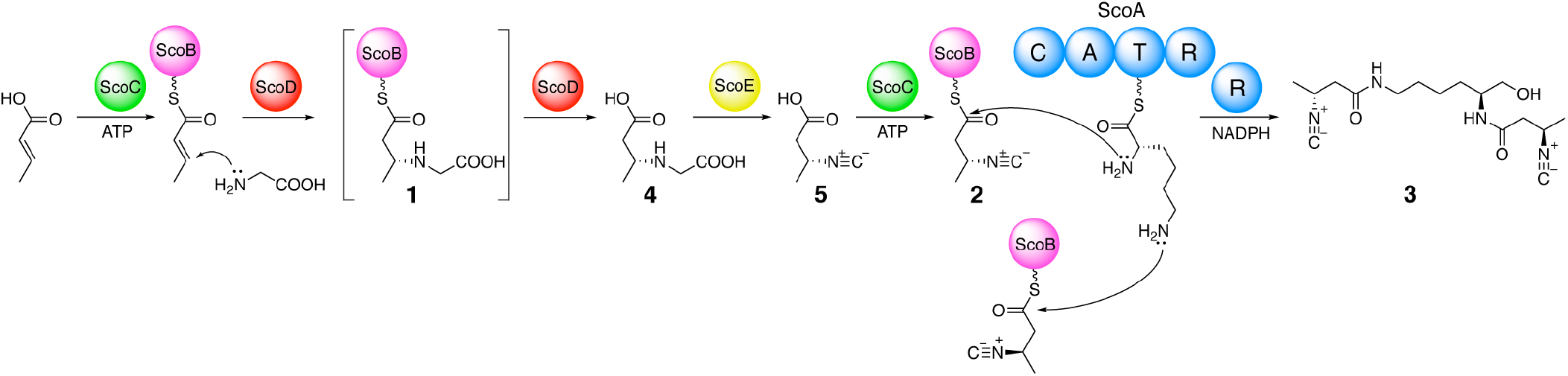
Schematic of isonitrile lipopeptide biosynthetic pathway. This revised pathway shows that ScoE utilizes a free acid substrate, **4**, and after isonitrile formation, the product **5** is reactivated and loaded onto ScoB by ScoC.

We previously proposed that ScoE functions on an ACP-bound intermediate because biochemical analysis of pathway enzymes showed that the formation of **1** requires ScoB, and the action of the NRPS, ScoA, also requires a ScoB-bound substrate. The proposed pathway thus accounts for the necessity and sufficiency of these five core biosynthetic enzymes for INLP synthesis (Figure 1). However the recent biochemical and structural characterization of three ScoD homologues indicated that these thioesterases have dual functions with enzymatic hydrolysis occurring immediately after the Michael addition, both steps mediated by a single Gly residue in the active site.^10–12^ These results raised the question of what the true substrate of ScoE is. We thus initiated an effort to reconstitute the *in vitro* activity of ScoE using various substrates that were chemically or chemoenzymatically synthesized. In particular, we chemically synthesized (*R*)-3-((carboxymethyl)amino)butanoic acid (CABA) **(4)** and CABA-CoA that was used alone or with the phosphopantetheinyl transferase from *Bacillus subtilis* (Sfp) to form CABA-ScoB **(1)** (**Figure S2-5**). Initial attempts to reconstitute the activity of ScoE were unsuccessful regardless of the substrate used.

Meanwhile, we obtained an X-ray crystal structure of ScoE to 1.8 Å-resolution (Figure 2; **SI Table 1**). The structure of ScoE is similar to the TauD family of non-heme iron(II) enzymes with root mean square deviation (RMSD) values of 2.1-2.7 Å (for C. atoms) to structurally characterized TauD enzymes, although the sequence identity is only 20-26% to these same enzymes.^13^ The 2-His-1-Asp facial triad (H132, D134, and H295) and an Arg (R310) within the substrate binding site are conserved with TauD family members. The ScoE substrate and alpha-ketoglutarate (αKG) binding sites are clearly visible and contain four exogenous ligands (Figure 2, A and C). Particularly, a Zn(II) is bound by the 2-His-1-Asp facial triad (H132, D134, and H295). The fourth coordination position on the Zn(II) is occupied by an acetate molecule that is acquired from the crystallization condition and is bound analogously to αKG in the structure of TauD (Figure 2, C and D).^14,15^ In the substrate binding pocket, a Cl^-^ is bound in a similar position to the sulfonate moiety of taurine in a substrate-bound structure of TauD (Figure 2, C and D).^14,15^ Surprisingly, a molecule of choline is found with the trimethylamine moiety oriented towards Cl^-^ (Figure 2C).

**Figure 2.**
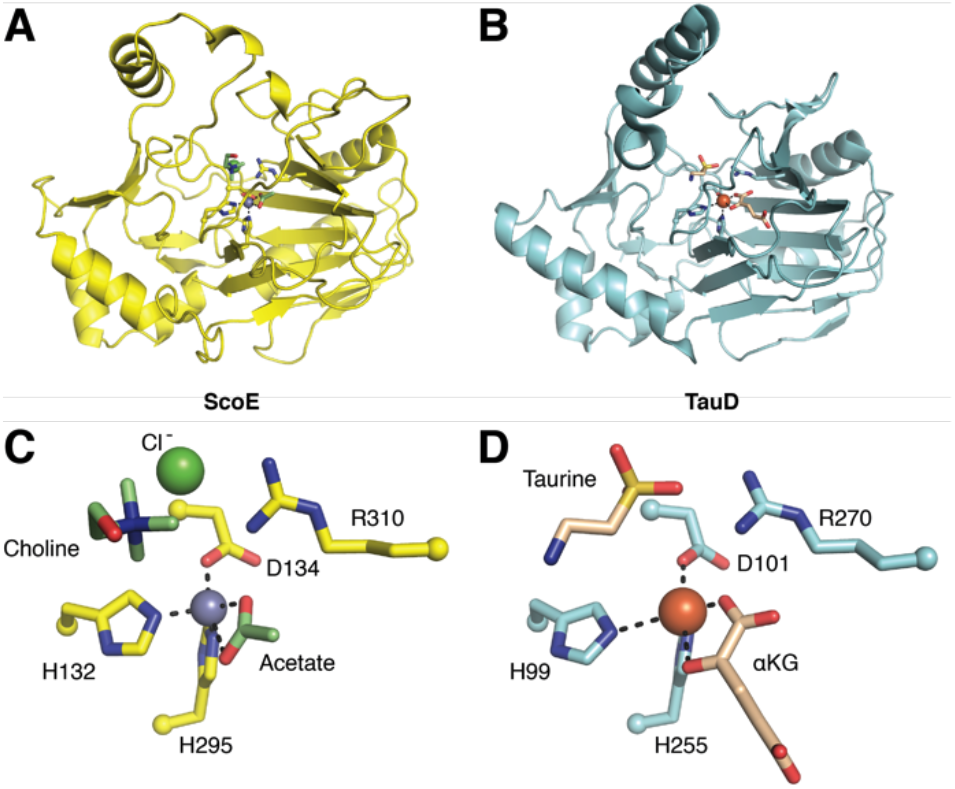
Structure of ScoE compared to TauD (PDB 1OS7).^15^ A) Overall structure of a ScoE protomer shown in yellow ribbon representation. The metal-coordinating 2-His-1-Asp facial triad and a conserved active site Arg are shown in yellow ball-and-stick representation. Acetate and choline ligands within the active site are shown in green ball-and-stick. Only one of the two observed conformations of choline is shown for clarity. Zn(II) and Cl^-^ are shown as green and gray spheres, respectively. B) Overall structure of a TauD protomer shown in teal ribbon representation and oriented similarly to ScoE in A). The metal-coordinating 2-His-1-Asp facial triad and a conserved active site Arg are shown in teal ball-and-stick representation. Taurine and αKG within the active site are shown in tan ball-and-stick. Fe(II) is shown as an orange sphere. C) Zoomed in view of the active site of ScoE. Composite omit density is shown in **Figure S7**. D) Active site of TauD displayed similarly to C).

Since choline and Zn(II) were identified within the crystal structure of ScoE but absent from the crystallization condition, we hypothesized that choline and Zn(II) were co-purified with ScoE during protein purification and they could potentially interfere with substrate binding and Fe(II) reconstitution of the holo-enzyme. To mitigate the problem of unwanted choline that may be abundant in LB media used for protein purification,^16^ we then used M9 defined medium for future purifications of ScoE (**Figure S6**). After affinity chromatography, ScoE was dialyzed in a buffer containing 1 mM EDTA to remove Zn(II). This apoprotein was reconstituted immediately before each assay with fresh Fe(II) and αKG to form the holo-protein.

We then incubated holo-ScoE with αKG, ascorbate, and either CABA-ScoB **(1)**, CABA **(4)**, or CABA-CoA. Upon incubation of ScoE with **4** and subsequent LC-HRMS analysis, we observed a mass spectrum associated with the formation of (*R*)-3-isocyanobutanoic acid **(5)** (Figure 3A; **S8**). This product was not observed when either **4**, αKG, or ScoE were omitted, or when boiled ScoE was added to the reaction. To further confirm this result, we synthesized (*R*)-3-((carboxymethyl)amino)butanoic-5-^13^C-acid as a substrate (**Figure S9**). When ScoE was incubated with this labeled substrate, the expected mass spectral shift of the product was observed (**Figure S8**). This mass spectrum was only observed when all necessary components of the ScoE reaction were included and only when the labeled substrate was utilized. Finally, we also confirmed the identity of the product from the enzymatic reaction by comparing to a chemically synthesized standard (Figure 3A). No activity of ScoE was observed when **1** or CABA-CoA was used as a substrate.

**Figure 3.**
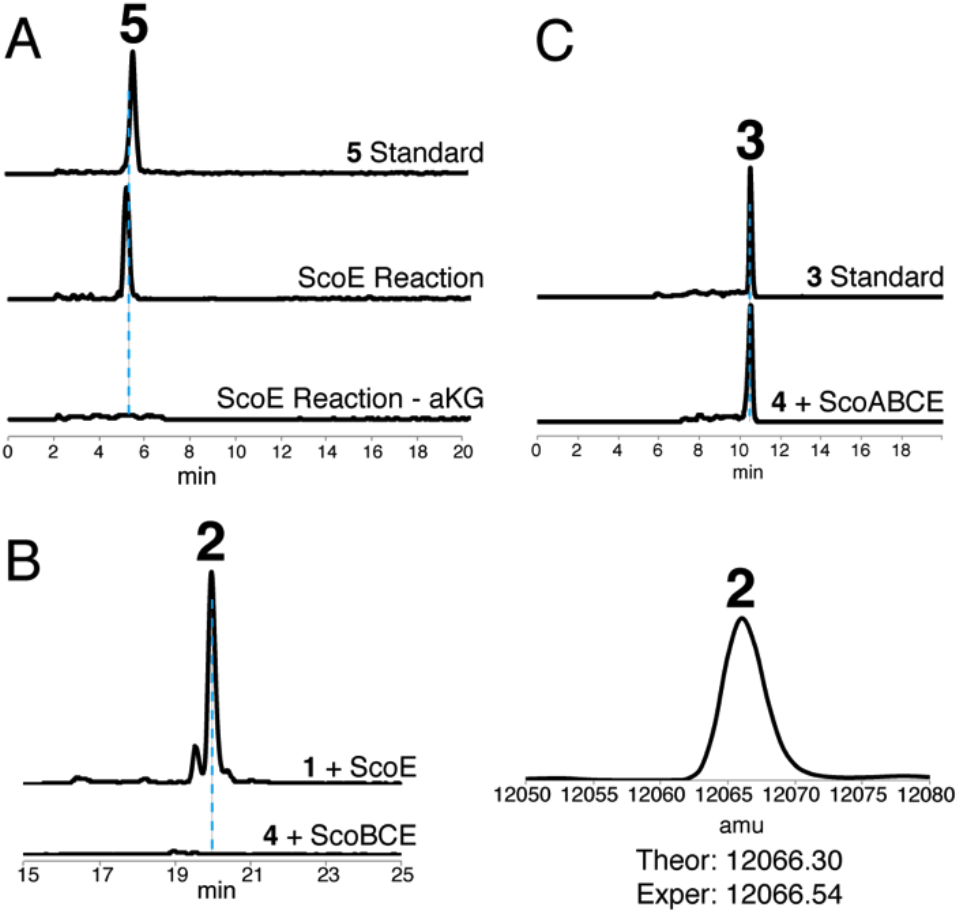
*In vitro* characterization of ScoE. A) Extracted ion chromatograms showing the conversion of **4** to **5** catalyzed by ScoE. For simplicity, only the no αKG assay is shown as a representative negative control. The calculated mass of 5 with a 10-ppm mass error tolerance was used. B) Extracted ion chromatograms showing the production of **2**. Bottom trace displays the assay with **1** and ScoE. Top trace shows the coupled reaction containing ScoE, ScoB, ScoC and **4**. The calculated mass of **2** with a 10-ppm mass error tolerance was used. The deconvoluted mass spectrum of **2** is displayed on the right. C) Extracted ion chromatograms showing the formation of **3** in the total enzymatic synthesis using ScoA, ScoB, ScoC, ScoE and **4**. The calculated mass of **3** with a 10-ppm mass error tolerance was used.

Our *in vitro* biochemical analysis provided direct evidence for a second mechanism of isonitrile formation by a non-heme iron(II) and αKG dependent oxidase/decarboxylase. We propose that ScoE functions similarly to TauD and TfdA,^14,17^ utilizing an enzyme-bound iron-oxo species for oxidation of **4** which likely goes through two sequential steps with an imine intermediate (**Figure S10**). The position of the choline hydroxyl group and Cl^-^ in our structure of ScoE may indicate the binding mode of the two carboxylate groups of **4**. However, a high-resolution structure of ScoE with substrate bound is necessary to determine the precise substrate binding arrangement and provide insight in substrate activation, which is currently under way.

The reconstitution of ScoE activity using the free acid substrate, **4**, raised additional questions about the function of enzymes in INLP biosynthesis. We have previously shown that the NRPS, ScoA, requires a ScoB-bound substrate for the subsequent amide bond forming condensation reactions. Our *in vitro* ScoE system however utilized only **4**, which after forming isonitrile product **5**, would need to be activated and loaded onto ScoB again for ScoA to function (Figure 1). We then tested whether the acyl-ACP ligase, ScoC, could function again after ScoE to activate **5**. We first conducted the ScoE *in vitro* reaction and after a short incubation period, added ScoB, ScoC, and ATP. The LC-HRMS analysis showed a strong signal for the formation of **2**, which was absent when **1** was directly used as a substrate for the reaction of ScoE (Figure 3B). This result indicated that ScoC functions twice in the pathway, to first activate and load crotonic acid, and subsequently to activate and load **5** onto ScoB to provide a preferred isonitrile substrate for ScoA (Figure 1). To further support the proposed functions of enzymes in INLP biosynthesis, we next performed an *in vitro* total enzymatic synthesis of INLP using purified enzymes including ScoA, ScoB, ScoC and ScoE. After incubating enzymes with **4**, ATP, Lys, and NADPH, the expected product **3** was successfully produced as compared to a chemical standard based on LC-HRMS analysis (Figure 3C; **S11**).

In summary, with the aid of a high-resolution crystal structure, we were able to circumnavigate inhibitory circumstances and biochemically reconstitute the activity of ScoE, a non-heme iron(II)-dependent enzyme, for isonitrile synthesis for the first time. We demonstrated that ScoE catalyzes the formation of isonitrile via the oxidative decarboxylation of the free acid substrate, **4**. We also provided evidence for a revised pathway for INLP synthesis with the second role of a promiscuous acyl-ACP ligase, ScoC, which activates the isonitrile product of ScoE before the NRPS-promoted INLP formation. This revision is expected to be applicable to other homologous INLP biosynthetic pathways found in Actinobacteria (**Figure S12**). This work paves the way to elucidate the enigmatic enzymatic mechanism for isonitrile formation using a non-heme iron(II)-dependent enzyme, of which homologues are conserved and critical for the virulence of pathogenic mycobacteria, including *M. tuberculosis*.

## Experimental Section

**Detailed experimental methods can be found in the Supporting Information.**

## Acknowledgements

N.C.H. is supported by the National Science Foundation Graduate Research Fellowship Program and D.A.B. by a National Institutes of Health (NIH) Molecular Biophysics Training Grant T32 GM008313. This research was financially supported by grants to W.Z. from the NIH (DP2AT009148), Alfred P. Sloan Foundation, and the Chan Zuckerberg Biohub Investigator Program. C.L.D. is a Howard Hughes Medical Institute Investigator. This work used Northeastern Collaborative Access Team beamlines (GM103403) and a Pilatus 6M detector (RR029205) at the Advanced Photon Source (DE-AC02-06CH11357).

